# Ruling the Roost: Avian Species Reclaim Urban Habitat During India’s COVID Lockdown

**DOI:** 10.1101/2020.12.15.422890

**Authors:** Raahil Madhok, Sumeet Gulati

**Affiliations:** Wildlife and Conservation Economics Lab, Department of Food and Resource Economics, University of British Columbia

**Keywords:** COVID-19, biodiversity, conservation, human-avian interactions, India, wildlife, coexistence

## Abstract

As we retreated to our dwellings in the “anthropause” of spring 2020, did other species return to our urban centres? We leverage an increase in balcony birdwatching, a million eBird entries, and difference-in-difference techniques to test if avian species richness rose during India’s COVID lockdown. We find that birdwatchers in India’s 20 most populous cities observed 8-17% more species during the lockdown. Most additional observations occurred after a two-week lag, signaling greater abundance instead of improved detection. More frequent appearances of at-risk, rare, and common species were recorded, implying that making our cities more wildlife friendly can protect threatened species in addition to urban specialists. Our contributions are: 1) to isolate and estimate a causal impact of reducing human activity on avian diversity, 2) to improve the external validity of this literature in rapidly urbanizing bio-diverse developing countries, and, 3) to illustrate a method separating abundance from detection in observational avian surveys.

## 1 Introduction

On March 27th, 2020, three days after India’s abrupt cessation of human activity, a reporter posted a video of a Sambar deer crossing a road in Chandigarh—among India’s most population dense cities^1^. Two days later, in the same city, a leopard was seen and tranquilized^2^. Similar sightings were reported elsewhere; a Nilgai walked along a usually bustling street in a Delhi suburb, a critically endangered Malabar Civet wandered alongside traffic in Kozikhode, Kerala, and flamingos returned in unusually large numbers to the Mumbai wetlands^3^. These sightings were not unique to India. As countries reined in human activity in response to COVID-19, animals were reported in previously busy locations all over the world^4^. Were these sightings chance events, or a systematic increase in animal presence in previously forbidding urban areas? The cascade of global COVID lockdowns provides a unique opportunity to shed light on this^5^.

We use citizen science data from before and during the lockdown to evaluate how the “anthropause” (a term coined by Rutz et al. (2020)) affects observed avian diversity in India. We also investigate if this change in observed avian diversity derives from a change in abundance, or from improved detection. Finally, we employ a species level analysis to provide a tangible understanding of the birds repopulating the cities in our results.

We study birds because they are a good proxy for overall wildlife, are easily observable, sensitive to environmental change, and comprise a broad taxonomy (Xu et al., 2018). We use over one million bird sightings from eBird^6^ to test whether observed species richness in India’s 20 most populous cities rose during the COVID-19 lockdown. As wildlife surveys stalled under lockdown^7^, eBird usage—with broad geographic and temporal coverage before COVID—soared. This allows us to present a robust and comprehensive analysis of our question.

India is among the best settings to learn about human-wildlife interactions. It juxtaposes a backdrop of unrivaled biodiversity, home to 12% of the world’s bird species (Jayadevan et al., 2016), against dense urban centres that are particularly forbidding for wildlife. 21 out of the world’s 30 most polluted cities (IQAir, 2020), and 4 out of the 10 most traffic congested cities (Index, 2020) are in India. These cities also have unhealthy noise levels^8^. In April 2020, as the world unleashed sweeping stay-at-home orders, India’s lockdown was among the most stringent (Hale et al., 2020) and featured a deep and sudden reduction in human activity (see section S1.1 for lockdown timeline).

Our main empirical challenge is to find a credible counterfactual. How would avian diversity have evolved in the absence of a lockdown? This is complicated because an annual out-migration of several species coincides with the lockdown period. We address this with a difference in difference (DD) technique (Angrist and Pischke, 2008) using the same 48 day period in 2019 as a counterfactual. We also restrict our sample to birdwatchers with similar skills, and make comparisons within the same hour-of-day, and within trips of the same type, to mitigate selection bias.

Our next challenge is to determine if changes in observed species diversity represent changes in abundance or improvements in detection—a challenge relevant to all those using observational surveys. We use a dynamic DD technique to estimate lockdown impacts over time. As changes in human activity, and associated reductions in noise and pollution occur almost immediately after lockdown, any sizeable lag suggests changes in avian presence rather than improved detection underlie our results.

Our analysis makes three important contributions. First, we estimate the causal impact of reducing human activity on avian diversity. We share this contribution with a nascent literature studying the impact of COVID lockdowns on wildlife elsewhere^9^. The related landscape ecology literature evaluates species diversity along an urban gradient (see Chace and Walsh (2006) for a review). Such analyses confound changes in human activity with changes in habitat^10^. In contrast, we compare species diversity within the same urban habitat, only human activity varies.

Second, we present robust estimates derived from a very large sample collected over a geographically and culturally diverse region, improving the external validity of our study to other developing countries where urbanization is accelerating and large swathes of species are imperiled (Newbold et al., 2016; Myers et al., 2000; Jenkins et al., 2013). So far, studies estimating the impact of COVID lockdowns on animal presence/behaviour use small samples in distinct study sites from the developed world. For example, Manenti et al. (2020) surveys water birds at an artificial lake during Italy’s COVID-19 lockdown and find higher species richness compared to 2019. Derryberry et al. (2020) find birds increasing their acoustic distance during San Francisco’s lockdown, and possibly improving breeding success^11^. In contrast, our estimates represent a statistically robust average impact of reduced human activity on avian diversity for 20 large cities across India.

Third, we illustrate a novel way to separate abundance from detection in observational surveys. Technology-based methods objectively capture animal *presence*, but observational surveys typically capture species *detection*^12^. Human observation is the primary method to collect data for avian surveys—whether via systematic surveys or by citizen science. Through our dynamic DD method, we formally investigate whether changes in species diversity represent changes in presence, or mere improvements in observer cognition from reduced noise and visual pollution.

## 2 Methods

### 2.1 Selecting a Counterfactual

An eBird ‘trip’ consists of a species checklist, information about the data collection process, and a ‘protocol’: stationary, or travelling (see section S1.2 and Table S1). To estimate the causal impact India’s lockdown on avian diversity, we need to compare species richness per trip during lockdown with a counterfactual period where lockdown never occurred. The weeks prior to lockdown is insufficient because the available species pool was different. Migratory species arrive during India’s warm winter and depart in Spring, coinciding with the lockdown period. A simple pre-post comparison would thus conflate the impact of lockdown with that of species migration.

We instead choose the same period in 2019 as the counterfactual. The assumption is that, had the lockdown not occurred, species richness would have evolved as it did during the same 48 day window in 2019 (known as the parallel trend assumption). The difference in species richness per trip before and after the fourth Wednesday in March 2019 captures species migration, and we subtract this from the pre-post difference in 2020. Any remaining “difference in difference^13^” will then be devoid of the migration bias.

### 2.2 Mitigating Selection Bias

Whereas our counterfactual accounts for species migration, it does not account for the changing nature of birdwatching after March 25th 2020. Figure 1 illustrates three such selection biases. Immediately following lockdown, there is a near-quadrupling of stationary users (Fig. 1A), and the number of trips they report^14^ (Fig. 1B), consistent with reports of surges in balcony birding in isolation^15^. To exclude new sign-ups, we first pre-process the data to improve representativity (see section S1.3) and then impose a ‘participation constraint’ to compare checklists from a constant user base with comparable skills (see section S1.4 and Table S3). In our preferred sample, users recording at least two trips in the 24 days before and after lockdown—called “consistent users”—are selected. Another bias arises from changing trip protocols. Fig. 1C shows substantially higher (collective) species richness on traveling trips pre-lockdown compared to stationary trips, because users can venture deeper into bird habitats. This trend reverses post lockdown. Without accounting for the trend reversal, species richness per trip *falls* during lockdown in Bangalore, Chennai, and Kolkata (driven by plummeting travelling trips), but *increases* within stationary trips (Figure 2). Therefore, we make all pre-post comparisons within trips of the same type to remove the protocol bias.

**Figure 1:**
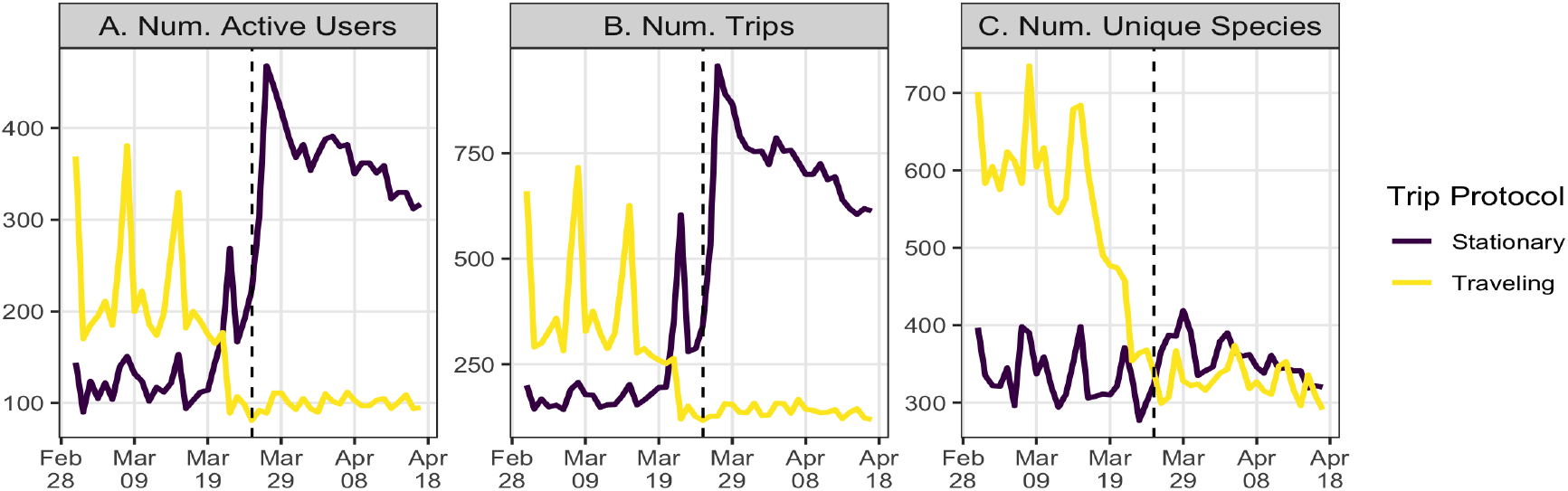
Daily Birdwatching Activity in India (March & April 2020, all cities) Note: Data include all trips from March 1st to April 17th, 2020 after applying the pre-processing steps described in the Section S1.3. Panel A plots the number of unique user ID’s active each day. Panel B plots the total number of daily trips. Panel C plots the daily number of unique species across all users and trips. The vertical dashed line is March 25th, 2020, the day lockdown was implemented.

Two additional sources of selection bias arise from the changing schedule of birdwatching during lockdown (figure S2.1) and rural-urban differences (discussed in section S1.4). We also control for a range of climatic and behavioural variables—including weather and trip duration—that change during our study window and also impact species diversity (section S1.5 and Table S4).

**Figure 2:**
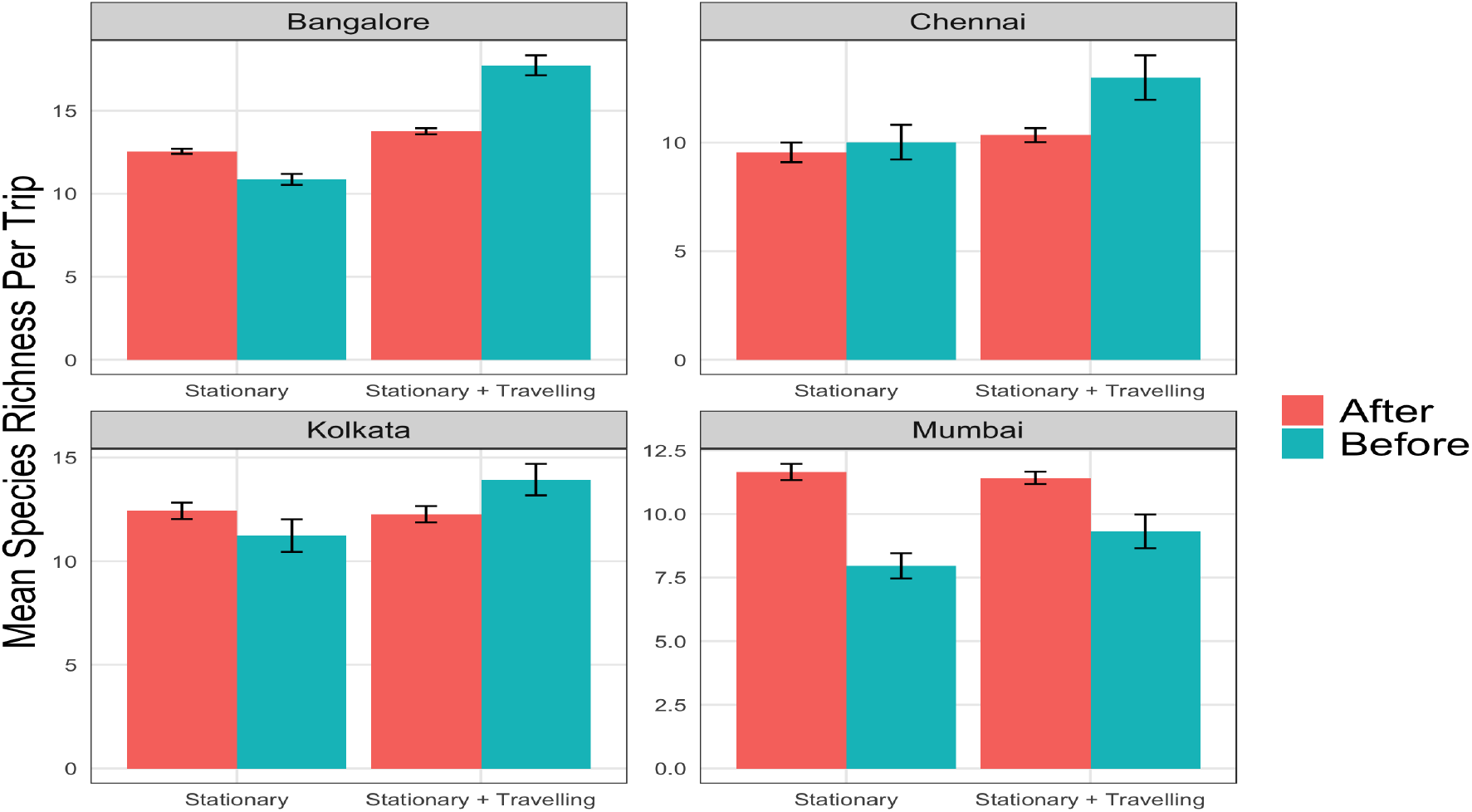
Species Richness per Trip in Four Cities (2020) Note: Data consist of trips taken by observers who recorded at least two trips in each of the 24 days before and after lockdown. Observations from Janta Curfew (March 22th) are dropped. Red bars are post-lockdown trips and blue bars are pre-lockdown. “Stationary + Travelling” pools together stationary and travelling trips. Black lines show the standard deviation of the mean.

### 2.3 Difference-in-Difference Model

Combining our counterfactual with the procedure for mitigating selection bias, we proceed to estimate a DD specification. The goal is to compare changes in outcomes in a treatment group before and after a policy date (the first difference) with changes in outcomes over the same period in a counterfactual where the policy was never implemented (second difference). We estimate:

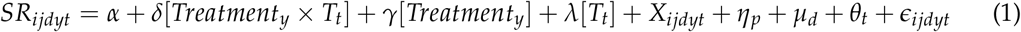

where *SR_ijdyt_* is species richness observed by user *i* on trip *j* in district *d* in year *y* and day-of-year *t*. *Treatment_y_* is a dummy for 2020. *T_t_* is a time dummy for the post period *t* ∈ [*Policy_t_*, *Policy_t_* + 24], where *Policy_t_* is the 4th Wednesday of March—the “policy date”. The pre-policy period is March 1st until *Policy_t_*, and the post-policy period is 24 days afterwards. *X_ijdyt_* is a vector of weather and trip covariates (section S1.4). *η_p_* is a protocol fixed effect that ensures all pre-post comparisons are made among trips of the same type. *μd* is a district fixed effect and ensures comparisons are made across trips within the same district (table S2). *θ_t_* is a set of temporal controls including hour fixed effects and weekend fixed effects. *δ* is our parameter of interest, denoting the causal impact of India’s COVID-19 lockdown on species richness, and is derived in section S1.6.

### 2.4 Dynamic Differences in Differences

The term *δ* in equation 1 captures the average impact of lockdown during 24 days of lockdown. We decompose this to parameterize marginal effects over time in order to investigate whether our estimates represent actual changes in species presence, or whether birds are just easier to observe in the absence of human activity. Since human activity stopped overnight, a lag between lockdown and higher species diversity suggests a gradual recovery of species. We estimate:

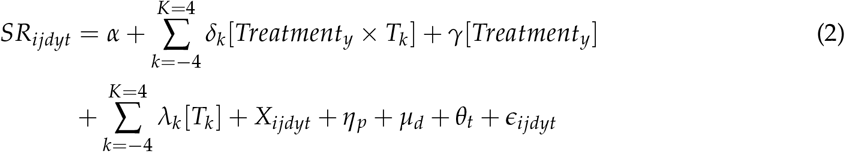

where *δk* is the DD estimate in time bin *k*. We use 8 bins with 6 days each^16^. *k* = (−6, 0] is omitted so that all estimates are relative to the six days before (and including the day of) lockdown. All other parameters are defined and interpreted in the same way as equation 1.

### 2.5 Marginal Species Identification

Since our outcome, *SR_ijdyt_*, measures the *number* of species per checklist, the DD estimate does not reveal *which* specific species are seen more or less often during lockdown. To investigate this, we first restrict our sample to that of the regression^17^, and calculate frequency distributions for individual species in 2019 and 2020 in Bangalore and Delhi—two cities with the most eBird activity. We then identify 19 “marginal” species (out of 203 total) in Bangalore and 23 (out of 138) in Delhi according to three criteria: 1) the average daily proportion of checklists reporting the species after the policy date is higher in 2020 than 2019; b) the average daily proportion of checklists reporting the species pre-lockdown in 2020 is no more than in 2019^18^; c) the species is observed on at least seven days in the post-lockdown period in 2020^19^.

Having established which species are observed more often during lockdown, we classify the rarity of these marginal species at a global and local level. For the global classification, we list the IUCN Red List category. Since this may misrepresent the local threat level, we classify a species as locally rare if their 2019 reporting frequency is in the bottom 25th percentile, a common threshold in the literature (Gaston, 1994).

## 3 Results

### 3.1 The Impacts of India’s Lockdown on Observed Avian Species Diversity

Figure 3 informally illustrates our DD method, without controls or fixed effects. Species richness on travelling trips drops after the policy date, but gradually increases among stationary trips ^20^. This corroborates a story of recovering species abundances (or diversity, see Discussion). Importantly, in the absence of lockdown (left of vertical dashed line), there was no systematic difference in species richness between 2020 and 2019, bolstering our choice of 2019 as the counterfactual.

**Figure 3:**
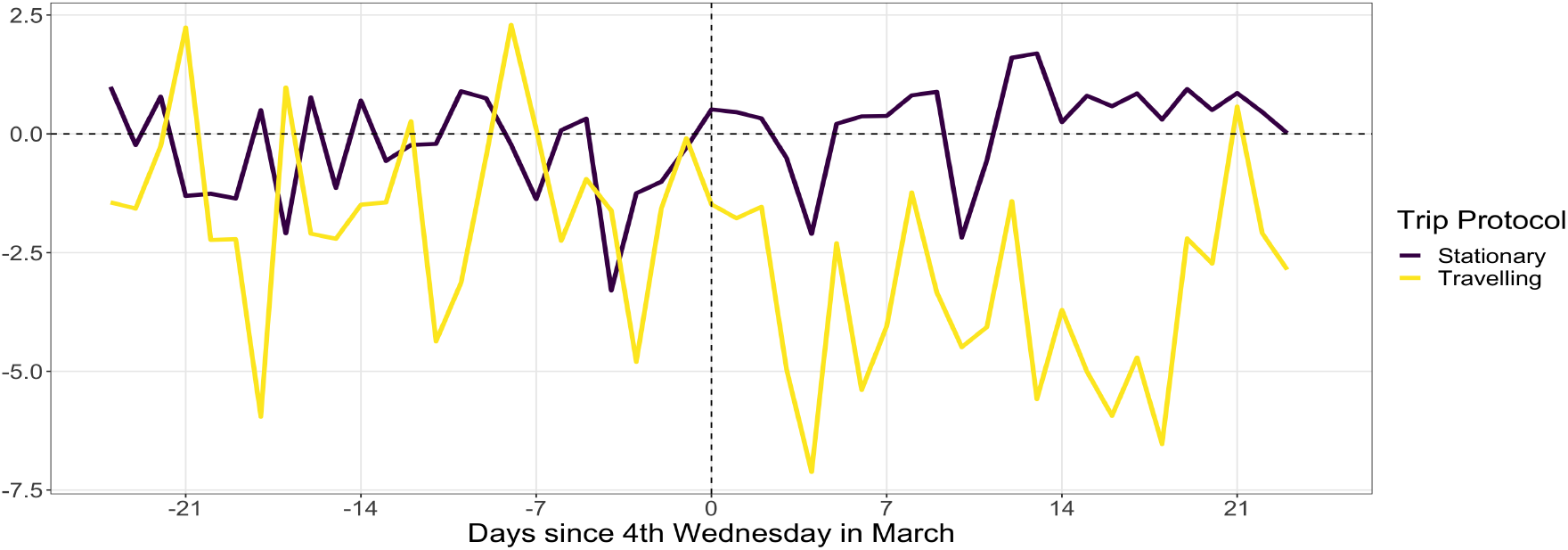
Daily Species Richness Per Trip Relative to 2019 in India Note: Data meets 2-trip participation constraint in both 2019 and 2020. The policy date (vertical line) is the 4th Wednesday of March, the date of lockdown announcement in 2020. Consistent users in 2019 need not be the same individuals as in 2020. Lines describe mean species richness per trip, coloured by protocol, across users on a given day in 2020 minus the value from the same day in 2019.

The formal DD estimates (equation 1) show robust evidence that a reduction in human activity increased observed avian diversity (Figure 4). The first coefficient—our preferred specification— shows India’s COVID-19 lockdown increased species richness by 1.23 species per trip (p*<*0.05) in the top 20 cities, equivalent to 8% of the pre-lockdown mean^21^.

**Figure 4:**
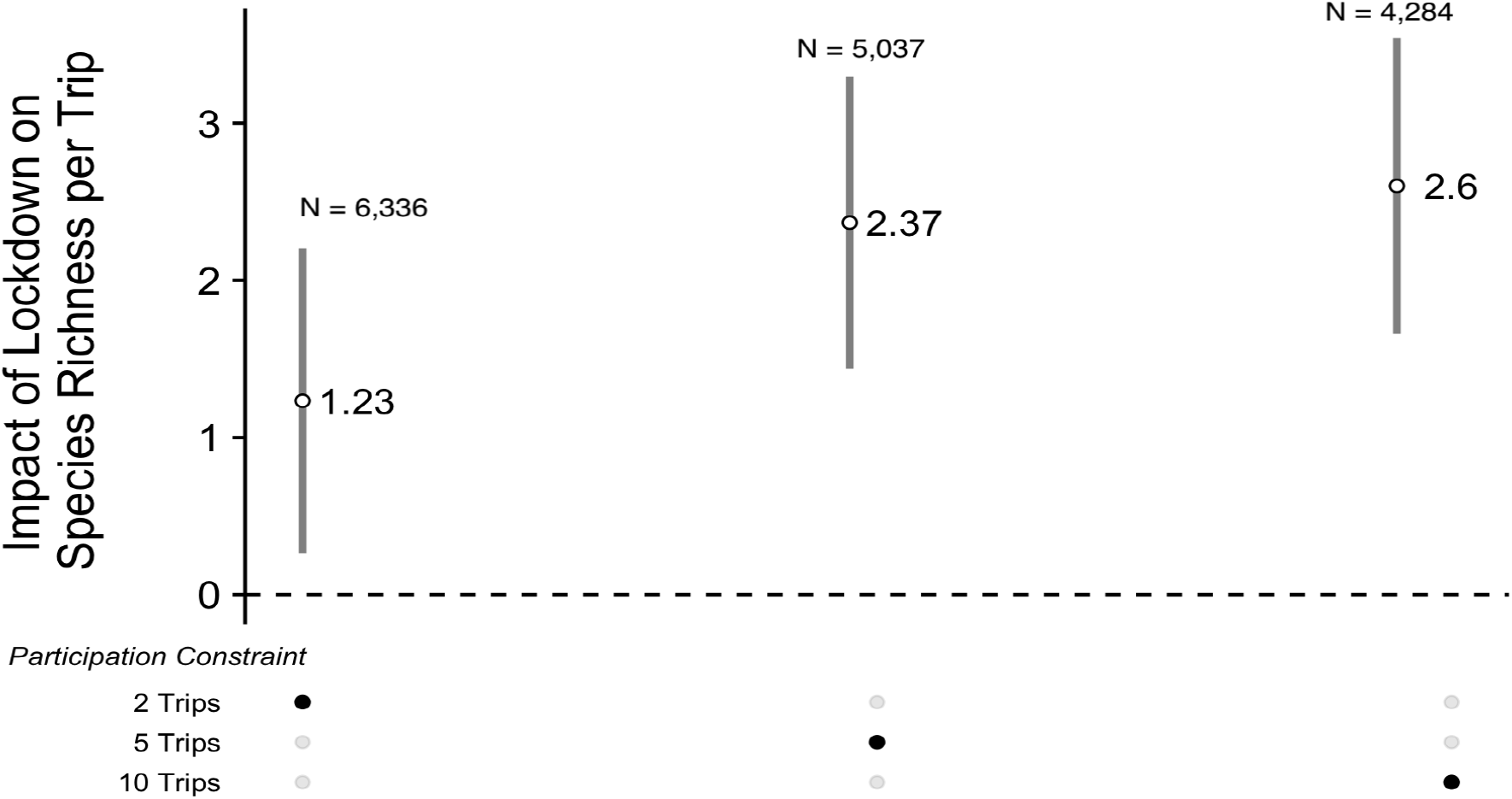
Difference in Difference Results Note: White circles describe the DD estimate from equation 1. Grey bars are 95% confidence intervals. The estimation sample is pre-processed, covers the top 20 cities, and imposes a participation constraint corresponding to the black button. Data from March 22nd, 2020 (the Janta Curfew) are dropped. All regressions include district, hour-of-day, and protocol fixed effects as well as controls for trip duration, rain, temperature, number of observers, distance to nearest birding hot-spot, and a weekend dummy. Standard errors are robust to heteroskedasticity.

There is a two-week lag (p*<*0.05) before an increase in species richness is detected (Figure 5). Pre-lockdown, there is no statistical difference in week-to-week species richness between 2020 and 2019, providing formal support for the parallel trend assumption. Our results are robust to a variety of alternative specifications (see section S1.7 and figure S3).

**Figure 5:**
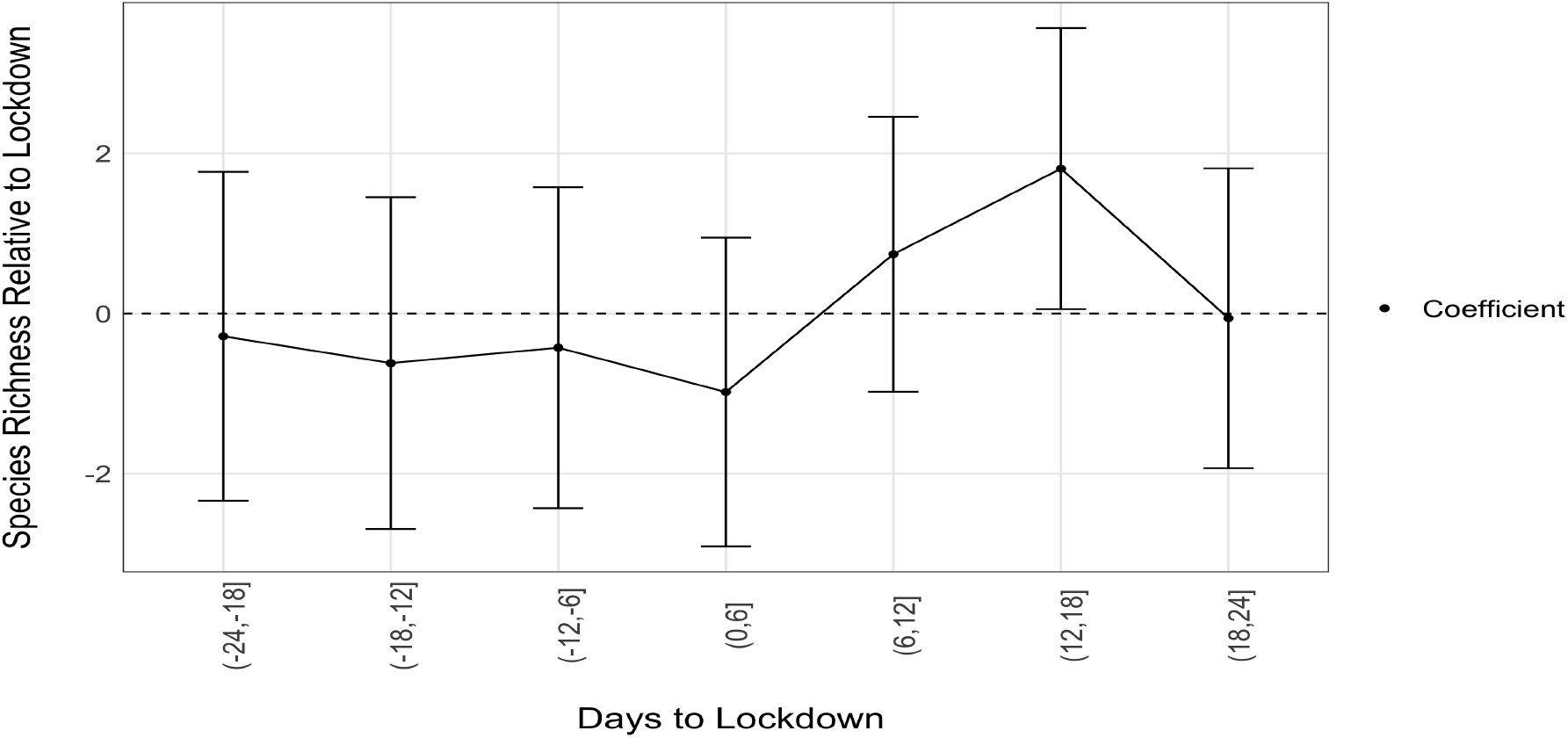
Dynamic Impact of Lockdown on Species Richness Note: The x-axis denotes 6-day time bins. Negative values denote days before lockdown. The coefficient is the DD estimate for the impact of lockdown on species richness during a given time bin relative to that same bin in 2019. The period (−6,0] is omitted so all coefficients are relative to the week before (and including the day of) lockdown. Bars are 95% confidence intervals. The estimation sample is pre-processed, cover the top 20 cities, and include users meeting the 2 trip participation constraint. Data from March 22nd, 2020 (the Janta Curfew) are dropped. Specification includes district, hour-of-day, and protocol fixed effects, and control for trip duration, rain, temperature, and a weekend dummy. Standard errors are robust to heteroskedasticity.

### 3.2 Which Species are Observed More often in Lockdown?

Figure 6 illustrates frequency distributions for four marginal species in Bangalore and Delhi (see Figure S5 for full set). In Bangalore, the Black-rumped Flameback Woodpecker is never reported in 2019, and never reported in the pre-period of 2020, but we find several reports of the species one week into lockdown. In Delhi, the Black-rumped Flameback is reported by a larger proportion of checklists in the pre-period in both years. In 2019, however, it is no longer observed during the post-period, but in 2020, continues to be reported throughout lockdown. A similar pattern is seen for the Large-billed Crow in Delhi: reported by a similar proportion of eBird checklists in the pre-period of 2019 and 2020, but only observed in the post period in 2020.

**Figure 6:**
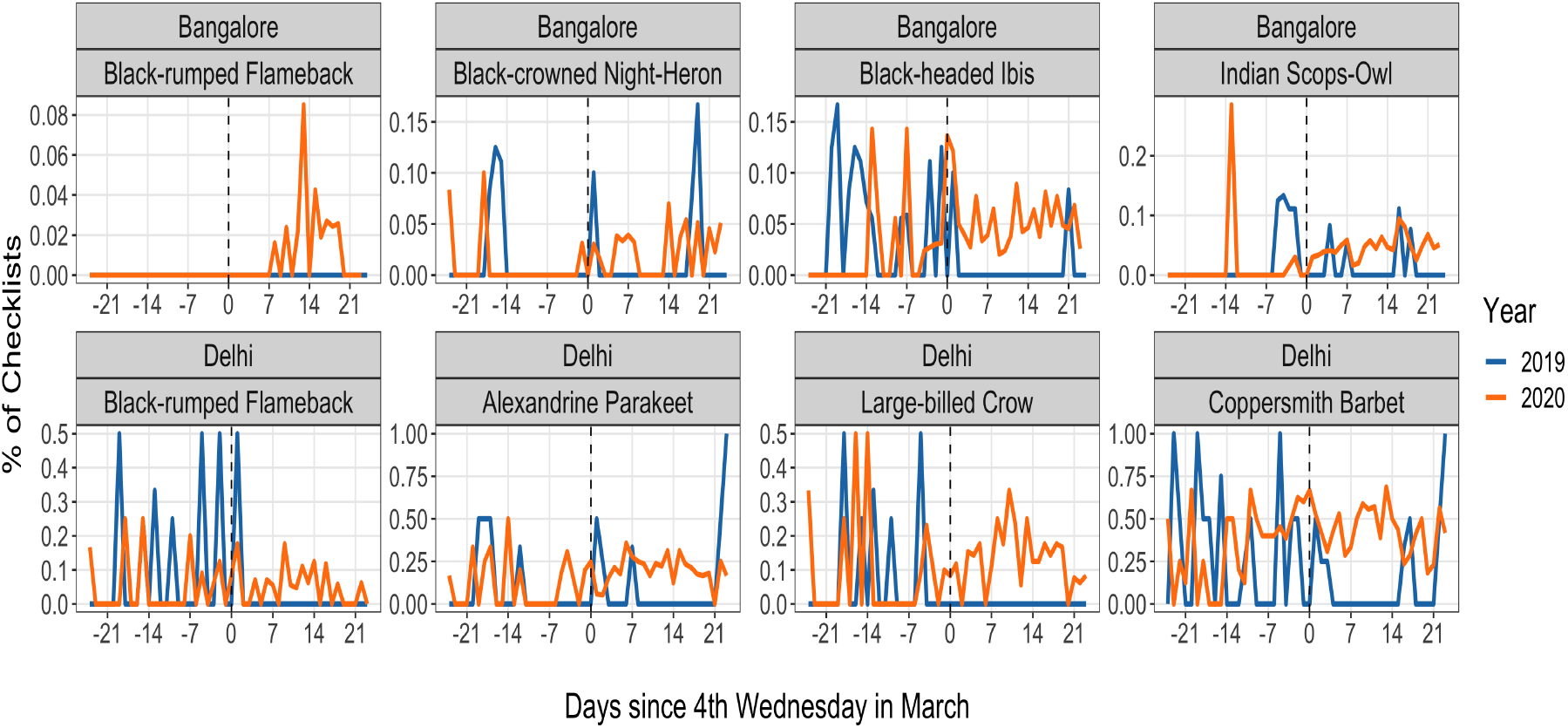
Marginal Species Distributions Note: Species Distributions of four marginal species in Delhi and Bangalore. Y-axis is the percentage of checklists reporting the species. X-axis is the number of days relative to the 4th Wednesday of March, which corresponds to the lockdown in 2020. The sample of checklists are stationary trips from the DD regression sample (2-trip participation contraint)

Among marginal species in Delhi and Bangalore, a handful are rare and the majority are common species detected more frequently (tables S5 and S6). The IUCN classification yields two globally near-threatened species: the Black-headed Ibis in Bangalore and the Alexandrine Parakeet in Delhi (distributions in Figure 6). The remaining marginal species in both cities are Least Concern. Our local rarity criteria classifies five (out of sixteen) marginal species in Bangalore as locally rare and four (out of nineteen) in Delhi.

## 4 Discussion

Our analysis presents a measurable change in the *viewing* of avian species. The cessation in human activity on March 25th 2020 was abrupt and strongly enforced. If the additional 1.23 species from our DD estimate were always present, but previously undetected because of distractions from human activity (e.g. traffic noise, and air pollution), then users should detect these species nearly immediately after lockdown when human activity stood still. Indeed, balcony bird-watching soars the next day. The fact that it takes two weeks to detect additional species suggests the abundance of incumbent species, or the emergence of species previously absent, gradually grew until the probability of detection was high enough two weeks after lockdown.

Our exploration of marginal species indicates both channels are active. Some species are seen frequently in Delhi and Bangalore before and after the 4th Wednesday of March in both years, but a greater proportion of checklists report them in the post lockdown period in 2020. Some are never seen in 2019 altogether, and only seen in 2020 during lockdown, exhibiting a “true” marginal observation.

We classify 26% and 17% of species seen more frequently during lockdown as locally rare in Bangalore and Delhi, respectively. One of these species in each city is globally near-threatened according to the IUCN. Rare species face greater extinction risk and are more sensitive to environmental changes (Gaston, 1994), making this analysis useful for allocating scarce conservation budgets. Our results suggest investment in making our cities more wildlife friendly can even protect some at-risk species, not just the urban specialists.

At least two environmental mechanisms can explain the increase in avian diversity and abundance from reduced human activity. First, the lockdown lowered **noise pollution**^22^. The abundance and occupancy of avian species are negatively impacted by noise pollution (Shannon et al., 2016), as elevated noise levels mask mating signals and defence mechanisms (Slabbekoorn, 2013)^23^. Since the lockdown reduced traffic and other anthropogenic noise, certain species may reoccupy the landscape in larger numbers^24^. However, lower noise pollution also increases the ability of observers to hear bird calls, an important clue for bird-watching.

The second possibility is a drop in **air pollution**. According to the 2019 World Air Quality Report for PM2.5^25^, 21 of the 30 most polluted global cities are in India. This includes 9 cities in our sample. During lockdown, these cities experienced unprecedented reductions in PM2.5 and other pollutants (Sharma et al., 2020; Mahato et al., 2020). Exposure to particulates can reduce species diversity (Sanderfoot and Holloway, 2017). Therefore, air quality improvements may underlie the higher species diversity we observe. However, lower air pollution, especially particulate pollution, also improves visibility, another important factor for species detection.

Our novel approach for disentangling the abundance and detection mechanisms suggest that landscape reoccupation from reduced noise and improved air quality during lockdown, and not improved detection, underlie our results. Regardless of the mechanism, there are important benefits from bird watching and engaging with nature (Bratman et al., 2019; Shanahan et al., 2016; Maldonado et al., 2018). A change in the viewing of birds thus represents a change in well-being for birdwatchers during an unprecedented time. Nevertheless, it behooves us to recognize that only those with the ability and resources to engage in leisure, in the face of the economic and humanitarian crisis precipitated by the pandemic, may be able to engage in this possibility.

## Footnotes

1 According to India’s census, Chandigarh’s population density of 9252 people per *km*^2^ is more than 24 fold its national density of 382 people per *km*^2^. The sighting was posted on twitter at https://twitter.com/ghazalimohammad/status/1243472152579342336?s=20, accessed, June 23rd, 2020.

2 https://www.tribuneindia.com/news/chandigarh/leopard-spotted-in-chandigarhs-sector-5-63171, accessed June 23rd, 2010.

3 https://www.cnn.com/travel/article/flamingos-mumbai-lockdown-scli-intl/index.html, accessed June 23rd, 2020.

4 Examples include: pumas in Santiago, Chile, dolphins in the Bosphorous, mountain goats in Wales, and deer in Nara, Japan, and Romford UK.

5 Other researchers (Rutz et al., 2020; Bates et al., 2020) also promote the value of the ‘Global Human Confinement Experiment’ (coined by Bates et al. (2020)) as an unprecedented opportunity to isolate the impact of human activity on animal presence.

6 The eBird Basic Dataset (EBD) (Cornell Lab of Ornithology, 2020) version ebd IN 201901 202004 relApr-2020, from the Cornell Lab of Ornithology (Sullivan et al., 2009).

7 Many animal surveys (including India’s avian surveys) were (and some still are) paused due to COVID-19 restrictions (Corlett et al., 2020). In India, the main recurring avian surveys are: the Asian Waterbird Census, conducted in wetlands only; the Great Backyard Count, conducted by volunteers on one weekend per year; and, the Common Monitoring Programme, assigning volunteers to a transect near their home in only one state.

8 See:https://www.livemint.com/news/india/which-is-india-s-noisiest-city-11581527059879.html, accessed June 3rd, 2020.

9 Our paper also relates to literature studying how lockdowns impacted other anthropogenic pressures on the environment such as pollution (see Dasgupta et al. (2002) for a review). The evidence is mixed (see Berman and Ebisu (2020), Persico and Johnson (2020), He et al. (2020), and Le et al. (2020)). In India, Mahato et al. (2020) find air quality improvements of up to 54% in Delhi. These studies provide valuable evidence on the human-environment nexus, and we add another, possibly less studied, dimension of species diversity.

10 For example, Xu et al. (2018) find avian species diversity declines by 26% as forestland transitioned into built-up areas in Shanghai. Verma and Murmu (2015) finds higher avian diversity in suburban relative to urban habitats in Jamshedpur, India. In both studies, the species declines can arise from a change in land-use, or associated human activity.

11 Indeed, in a separate survey of the Common Swift, (Manenti et al., 2020) found more eggs laid per female during lockdown.

12 Terrestrial species are increasingly monitored with autonomous cameras (Silveira et al., 2003), GPS collars, and radio collars (Cagnacci et al., 2010). Note that this is slowly changing as GPS transmitters become small and light and can be carried by birds (documented in the Movebank database, Wikelski M, Davidson SC, Kays R [2020]. Movebank: archive, analysis and sharing of animal movement data. Hosted by the Max Planck Institute of Animal Behavior (see http://www.movebank.org, accessed on August 9th, 2020).

13 A commonly used econometric method to estimate causal estimates in natural experiments. Please see Angrist and Pischke (2008) for a practitioners guide.

14 There is also an unsurprising reduction in users taking travelling trips. Although travelling is generally not permitted during lockdown, there are exceptions. These might be government officials (for example forest officials) or civilians walking within a large housing complex.

15 For a reference see https://www.nytimes.com/2020/05/29/science/bird-watching-coronavirus.html, accessed, June 24th, 2020.

16 We choose a 6 day window so that every bin has the same number of days (48 days / 8 bins = 6 days per bin).

17 We apply a 2-trip participation constraint across the top 20 cities. Then we choose stationary trips to make the 2019 and 2020 distributions comparable.

18 Because if true, the species is consistently more common in 2020, not only after lockdown.

19 Note that some species may fit most of our criteria, but only be observed one or two times post-lockdown. These will not be labelled ‘marginal’, but still contribute to the average increase in species diversity in our regression estimates.

20 Figure S2 shows the same illustration for number of users and trips. Stationary activity spikes and travelling activity drops after the 2020 policy date compared to 2019 due to mobility restrictions in the former but not latter

21 The other two coefficients show that tightening the participation constraint to five and ten trips increases the point estimates to 2.37 (15% increase) and 2.60 (17% increase), respectively, and improves precision. This is likely due to the more experienced sample. However, the number of selected users drops, and sample size reduces by 15% each time

22 A February 2020 report in LiveMint, an e-paper in India, reports that average noise levels recorded by monitors in residential areas of Mumbai, Delhi, Bengaluru, Kolkata, Chennai and Hyderabad were 65.4 db, 10 db higher than the maximum recommended by the Central Pollution Control Board. See: https://www.livemint.com/news/india/which-is-india-s-noisiest-city-11581527059879.html, accessed June 3rd, 2020.

23 Shannon et al. (2016) reviews papers analyzing the impact of noise on wildlife, finding a majority reporting altered vocal behaviour, reduced abundance in noisy habitats, changes in vigilance and foraging behaviour, and impacts on individual fitness and community structure.

24 Another paper isolates the impact of noise, separate from that of the other impacts of traffic and habitat finding moderate traffic noise reduces bird abundance by one-quarter, and brings about an almost complete avoidance by some species (McClure et al., 2013).

25 Accessed at https://www.iqair.com/world-most-polluted-cities/world-air-quality-report-2019-en.pdf, downloaded, June 4th, 2020.

## Acknowledgements

We thank participants of the UBC WCEL Seminar for helpful feedback. We thank Tatiana Zarate for suggesting to control for distance to hotspots, and we thank Patrick Baylis for recommending two of our robustness checks.

## Author Contributions

Raahil Madhok conceived the idea for this paper, downloaded eBird data and designed the initial analysis, implemented subsequent analyses, and created all table and figures. Sumeet Gulati influenced data analysis, influenced and guided presentation and formatting of tables and figures. Both authors wrote sections of the paper. Both authors reviewed the manuscript

