## Supplementary Materials for "Ruling the Roost: Avian Species Reclaim Urban Habitat During India’s COVID Lockdown"

December 15, 2020

*

### S1 Materials and Methods

#### S1.1 Study Context: The Response to SARS-COV2 in India

India's first case of COVID-19 (the disease associated with the virus SARS-COV-2) is confirmed in the state of Kerala on January 30th 2020[1]. During February and March, official case numbers slowly rise. On March 11th, the World Health Organization declares SARS-COV-2 a global pandemic. India's first official COVID-19 death is reported the next day. It occurs in the state of Karnataka, and in response the state's chief minister orders all cinema halls, schools, and pubs to close for one week. The central government suspends visas to India, ordering a mandatory quarantine for visitors from affected countries[2]. Over the next two days, there is a cascade of state-level closures of schools, malls, cinema halls and popular gathering spots.

As concern around the disease rises (Beyer et al., 2020), at 8 PM on March 19th, Prime Minister (PM) Narendra Modi requests a voluntary nationwide stay-at-home period (termed 'Janta Curfew') for 14 hours from 7 AM until 9 PM, March 22nd. That same day, his government also announces a week-long moratorium on international flights to India from March 21st at midnight[3]. The announcement of Janta Curfew creates widespread speculation of an impending full lockdown[4]. On March 22nd, Janta Curfew is observed nationwide, and the central government also announces the suspension of passenger services on the Indian Railways until March 31st. Two days later, on March 24th at 8PM, PM Modi addresses the country once again. This time he announces a mandatory 21 day nationwide stay-at-home order beginning midnight March 25th (within four hours of his announcement)[5]. This includes all forms of transportation—public, commercial, taxi, and private. Only essential goods can be transported. Individuals can only leave home for urgent medical care, or with government-approved curfew passes. On April 14th, the lockdown is extended until May 3rd, with some relaxation for some districts with low infection rates after April 20th[6]. Freight transport is allowed after April 20th, and some local businesses can open. Agricultural and aquaculture operations are allowed to resume, some public works are restarted. On the 1st of May the nationwide lockdown is extended until May 17th. And finally, on May 17th, the nationwide lockdown is extended until May 31st with some further relaxations nationwide.[7]

[1]This description is written after referencing the following sources. a) The Times of India. A timeline of COVID 19, and government action: https://www.timesnownews.com/india/article/covid-19-a-comprehensive-timeline-of-coronavirus-pandemic-in-india/579026, last accessed, Wed, June 3rd, 2020; b) Forbes News India has a timeline at: https://www.forbesindia.com/article/coronavirus/coronavirus-timeline-how-did-we-get-here/58515/1, last accessed, Wed, June 3rd, 2020; c) For an update on lockdown extensions, see: https://www.timesnownews.com/india/article/india-coronavirus-lockdown-extended-to-phase-4-here-is-looking-into-the-dates-of-all-the-lockdowns-uptil-now/585837, last accessed, Wed, June 3rd, 2020; d) A Wikipedia article on India's lockdown has additional sources supporting the timeline above: https://en.wikipedia.org/wiki/COVID-19_pandemic_lockdown_in_India, last accessed, Wed, June 3rd, 2020

[2]See https://www.cnbc.com/2020/03/12/coronavirus-india-suspends-most-visas-closes-land-border-with-myanmar.html

[3]See https://www.thehindu.com/news/national/narendra-modi-speech-live-updates-coronavirus/article31108793.ece

[4]The day before Janta Curfew, members of the press document a run on essentials, see https://www.thehindu.com/news/national/kerala/panic-buying-marks-janata-curfew-eve/article31132243.ece.

[5]See https://www.nytimes.com/2020/03/24/world/asia/india-coronavirus-lockdown.html

[6]A classification of zones with different levels of restrictions based on infection rates is announced on April 20th

[7]Some private offices open, with workers returning on staggered schedules. Malls, metro trains, schools, colleges, hotels, restaurants, cinema halls, shopping malls, gyms, swimming pools, and places of religious worship remain closed. Some long-distance passenger train and air travel are reintroduced.

#### S1.2 eBird Data

During a birdwatching session with eBird, which we call a trip, the app records several trip characteristics that are uploaded to a centralized database. A trip includes a species checklist, information about the data collection process, location and date identifiers, and a group code to identify users on the same trip. The checklist includes the common and scientific name and the count of each species observed. If unable to count the birds, the user can input an "X" to indicate species presence. The main information on data collection are: trip duration, whether the checklist includes all species or only highlights, trip GPS coordinates, a name for the location (optional), and the trip protocol.

We obtained the eBird Basic Dataset (EBD) product for India from the Cornell Lab of Ornithology (Cornell Lab of Ornithology, 2020; Sullivan et al., 2009) for March and April of 2019 and 2020. The sample frame contains 1,586,974 observations across 100,962 trips and 5,037 users. Table S1 shows the breakup of trip protocols. 92% of trips are stationary or travelling. Travelling protocols are specified when a user moves more than a slight distance and they must enter the distance travelled. The stationary protocol is used when the user remains in one place. There is some vagueness of definition. For example, if a user moves from one side to the other side of their balcony, this would be considered stationary. For more details see Chapter 2 of the eBird Best Practice Manual (Strimas-Mackey et al., 2020). The remainder of trips are mainly incidental—where bird-watching is not the primary activity—but also include at-sea trips (Pelagic Protocol), research trips (Nocturnal and Shorebird Survey), and historical entries. Less than 10% (n=9065) report only highlights, which we consider non-representative of local species diversity.

#### S1.3 Data Pre-Processing

We apply the following criteria to identify representative checklists: a) keep stationary and travelling trip protocols; b) keep trips where all species observed are reported; b) drop duplicates from group trips where a leader's list is copied to participants' accounts; c) keep trips with duration between 5 and 240 minutes (Callaghan et al., 2017); d) drop trips recording a single species.

After removing these non-representative trips, we overlay remaining trip GPS coordinates onto the 2011 Census district map obtained from `https://www.diva-gis.org/gdata`. We drop 109 trips that fall just outside the coastline and mainly describe boating trips. 87 of the remaining trips have missing district names. We assign to these the 2011 district names corresponding to the trip coordinates.

The final data include 1,080,034 observations across 68,000 trips and 3600 users. Their collective checklists include 1433 unique species. This "pre-processed" sample standardizes the representativeness of trips, in terms of describing species diversity in the area, but does not account for new sign-ups and other biases. We layer the location and participation constraints onto the pre-processed data to obtain the final analysis sample.

#### S1.4 Removing Selection Bias

We use information on the data collection process to account for simultaneously changing observer and trip characteristics. First, we impose a "participation constraint" to compare checklists from a constant user base. In our preferred sample, only those recording at least two trips in the 24 days before and after lockdown—called "consistent users"—are selected. Formally, users who have at least two trips logged for each period March 1st-24th, and March 25th-April 17th are included. This excludes new post-lockdown sign-ups, who due to their inexperience, might log fewer species per trip (Zhou et al., 2020). Table S3B shows that 103 users meet a participation constraint of having taken at least two trips in the 24 days before and after lockdown. The collective number of trips among them more than doubles after lockdown.

Second, we make pre-post comparisons only within trips of the same protocol. Formally, this is done by including fixed effects for trip protocol in our difference in difference regression. Otherwise, the steep reduction in post-lockdown traveling trips would imply the lockdown is associated with a reduction in species diversity (Figure 1 in main text). Similarly, we compare observations among trips taken *in the same hour of day* to ensure the changing structure of bird watching times does not bias our results. Figure S1 illustrates a higher proportion of post-lockdown trips recorded between 4am-7am, coinciding with the time when many species are more vocal (Kelling et al., 2015). We address this temporal bias by including time of day fixed effects in our regression.

Lastly, while human activity falls in sparsely populated districts, it is low to begin with. Thus, if we pool together countrywide observations, we aggregate a low effect in rural areas with a possibly large impact of activity reduction in densely populated areas. To highlight the impact of human activity, we restrict our sample to cities within the top 20 by population density (see Table S2 for a list).

#### S1.5 Covariate Construction

Our covariates describe environmental conditions and birdwatching effort. To measure changing environmental conditions, we collect high-resolution satellite data on rainfall and temperature. Satellite data represents a substantive improvement over ground monitor data due to the dense spatial and temporal coverage. Rainfall (in mm) is collected on a $0.1 \times 0.1$ degree grid from the NASA Global Precipitation Measurement Product (Huffman et al., 2019). Gridded temperature data (in degrees Celsius) is collected from from the ERA5 global atmospheric reanalysis product maintained by the Copernicus Climate Change Service (Copernuicus, 2020) [8]. For each of these datasets, we overlaid the 2011 census district map and extracted the mean over all grid cells within district boundaries to create a district-daily panel of rainfall and temperature.

We collect data on six variables to measure birdwatching effort: distance to nearest hotspot (in km), trip duration (in minutes), hour-of-day of trip, number of observers, trip protocol, and an indicator for whether the trip was taken on a weekday or weekend.

A hotspot within a city is a public birdwatching site suggested by eBird users and approved by an eBird administrator. Typically, a location such as ponds, parks, etc, where a greater diversity of birds are seen. Before the lockdown, users could access these hospots, but not afterwards. We obtained a list of GPS coordinates for all hotspots in India from the eBird API[9] as of July 2020. To compute distance to the nearest hotspot, we measure the Great Circle Distance (in km) from each trip's latitude-longitude coordinates to the nearest point from the hotspot list. To control for changes in time spent birding on each trip, we also control for trip duration. Table S4 demonstrates that users are approximately no further from hotspots than earlier. However, we find that trips are, on average, shorter after lockdown. For the remaining covariates, all variables

[8]We use the specific dataset called "Land Hourly at Single Levels"
[9]See www.confluence.cornell.edu/display/CLOISAPI/eBird-1.1-HotSpotsByRegion

are recorded through the app and directly reported in the EBD.

##### S1.6 Coefficients and Assumptions in Difference in Difference Estimation

To interpret the coefficients of equation 1 in the main text under the intuitive DD framework, note that there are four parameters ($\alpha$, $\delta$, $\gamma$, $\lambda$) and the conditional expectation of $SR_{ijdyt}$ can take on four values depending on $Treatment_t$ and $T_t$. We hold constant $X_{ijdyt}$ and omit it from the conditional expectation to simplify notation. The formulas for $\alpha$, $\gamma$ and $\lambda$ are:

$$\alpha = E[SR_{ijdt}|y = 2019, t < T_t] \quad (1)$$
$$\gamma = E[SR_{ijdt}|y = 2020, t < T_t] - \alpha \quad (2)$$
$$\lambda = E[SR_{ijdt}|y = 2019, t > T_t] - \alpha \quad (3)$$

Intuitively, $\alpha$ is the baseline, pre-policy mean species richness in 2019. $\gamma$ is the pre-policy difference in species richness between 2020 and 2019. $\lambda$ is the time trend in 2019 (which in our context captures species migration). Following this notation, the difference in difference we are interested in can be written as:

$$\{E[SR_{ijdt}|y = 2020, t > T_t] - E[SR_{ijdt}|y = 2020, t < T_t]\} \quad (4)$$
$$- \{E[SR_{ijdt}|y = 2019, t > T_t] - E[SR_{ijdt}|y = 2019, t < T_t]\}$$

which is exactly equivalent to $\delta$ in regression equation 1 of the main text, our coefficient of interest. To see complete derivations, see Chapter 5.2 of Angrist and Pischke (2008). To interpret $\delta$ as the *causal* impact of lockdown on species richness, the parallel trend assumption must hold; that if lockdown never occurred, species richness in 2020 would have evolved similarly to its 2019 trend. The lack of pre-trends in figure 3 of the main text suggests the assumption holds, and the pre-period null effects in the dynamic specification (figure 5 in main text) provides robust statistical support.

##### S1.7 Robustness of Results

Figure S3 shows results from a range of robustness checks that test the sensitivity of our results to alternative specifications. Estimating the DD equation without controls for distance to hotspot and number of observers leaves the result virtually unchanged (Column 1). In column 2, we test the importance of the location constraint by removing it altogether (but keeping the 2-trip participation constraint). The point estimate drops to 0.233 and becomes insignificant, confirming our conjecture that including all locations dampens the impact due to the addition of rural areas where human activity was unaffected by the lockdown.

We also investigate the concern that users with different abilities are pooled together when calculating the pre-post differences. Differences in fixed observer characteristics could potentially bias the final DD estimate. The threat from this bias is arguably low because the participation constraint already ensures users have a consistent level of experience and our covariates ensure similarity along several other dimensions. Nevertheless, we re-estimate the DD equation such that the pre-post difference in each year is computed *within* a user [10]. In column 3, the point estimate with a 2-trip participation constraint drops to 0.78 and is statistically significant at the 10% level. This attenuation could arise from insufficient variation to detect a strong effect since each calculation is made over a minimum of two observations. As noted in Angrist and Pischke (2008), coefficients from fixed effect models attenuate toward zero as more restrictive fixed effects are considered. In line with this, when the participation constraint is loosened to five trips (column 4), the point estimate becomes 1.09 and regains statistical significance ($p<0.05$), almost equivalent to our preferred specification.

We also test the robustness of our dynamic DD specification under the more restrictive 5-trip participation constraint (Figure S4). The increase in species richness in the second week of lockdown is once again observed. The magnitude is slightly larger and more precise than the main specification, perhaps because of the higher experience level of the selected users. An increase in the first week is observed as well. There is also an increasing (but mostly insignificant) trend pre-lockdown, possibly due to small sample bias induced by the low probability that users logging 5 trips before and after lockdown are active in a given week.

[10] Since the user base is difference in 2019 and 2020, we cannot add user fixed effects in our main DD equation. Instead, we estimate the pre-post difference in species richness with user fixed effects in each year separately, and subtract the coefficients manually to obtain the DD estimate.

### S2 Supplementary Tables and Figures

#### S2.1 Figures

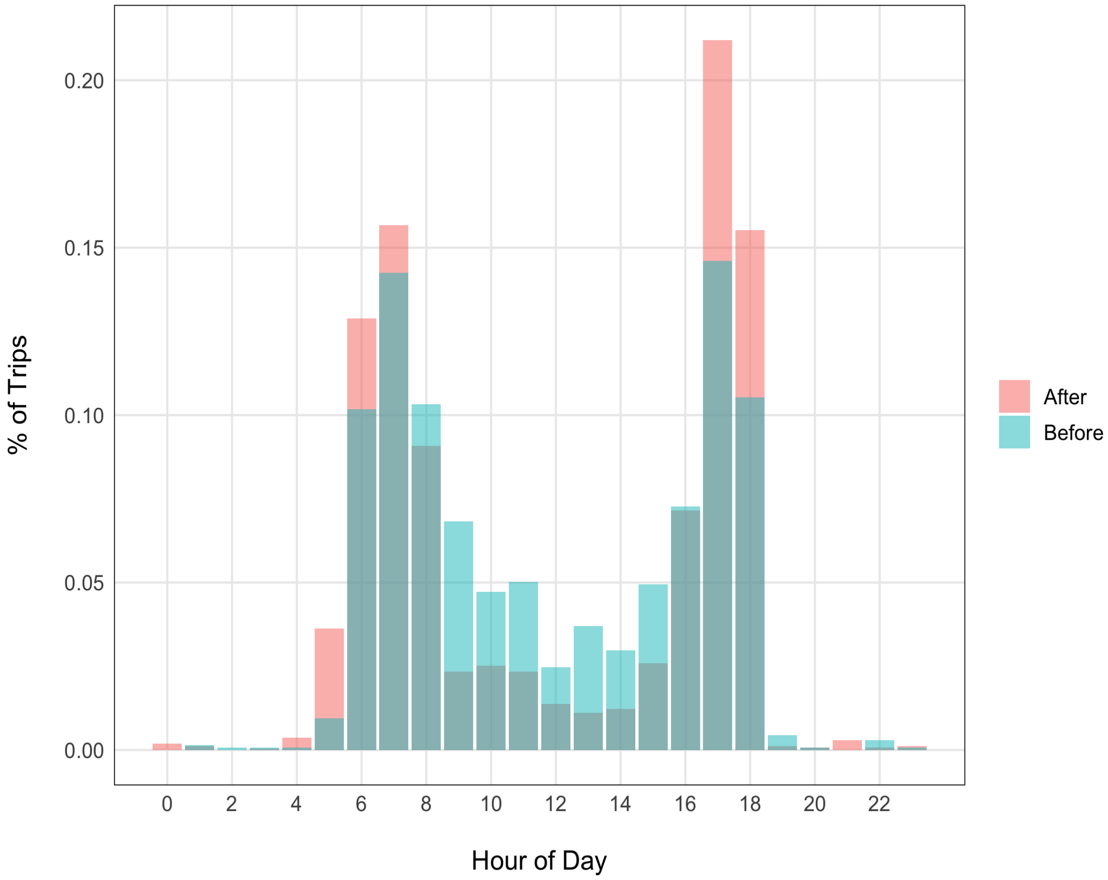

Figure S1: Hour-of-day Distribution of Trips (2020)

Note: Data consist of trips taken by observers in the top 20 cities by population density who recorded at least two trips in each of the 24 days before and after lockdown. Observations from Janta Curfew (March 22th) are dropped. Red bars describe the proportion of total post-lockdown trips recorded at each hour. Blue bars describe the same for pre-lockdown trips.

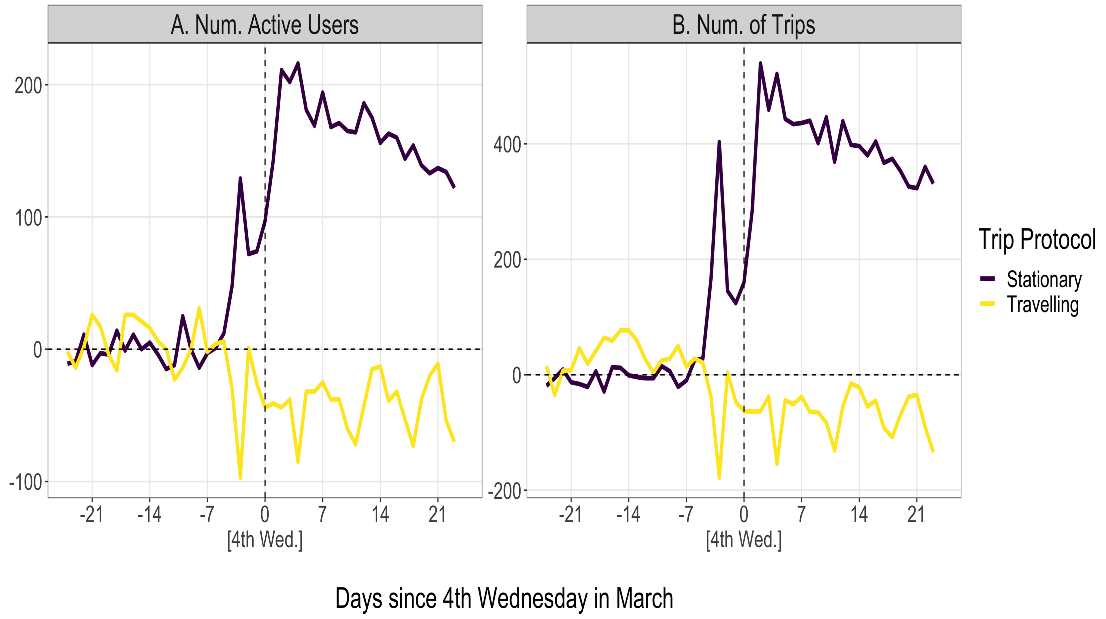

Figure S2: Daily Birdwatching Activity in India relative to 2019

Note: Data include trips from all users meeting the 2-trip participation constraint in both 2019 and 2020. Specifically, the reference period is set as the 4th Wednesday of March, corresponding to the date of lockdown announcement in 2020. Trips by consistent users in 2019 are appended to trips by consistent users in 2020, and these need not be the same individuals. Lines describe the number of active users (panel A) and total number of trips (panel B), coloured by protocol, on a given day in 2020 minus the value from the same day in 2019.

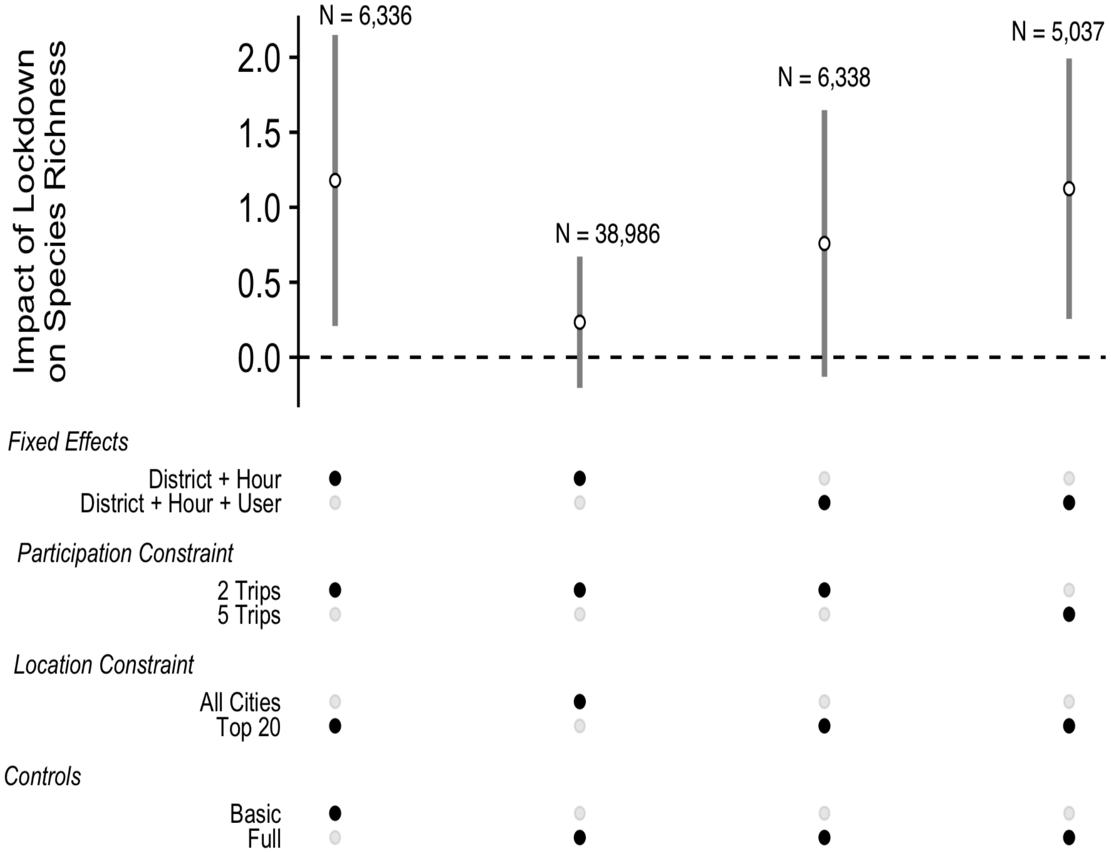

Figure S3: Robustness Checks

Note: White circles describe the DD estimate from equation 1. Grey bars are 95% confidence intervals. The estimation sample for all regressions is pre-processed for representativity, and include users meeting the participation and location constraint indicated by the black buttons. Data from March 22nd, 2020 (the Janta Curfew) are dropped. All regressions include district, hour-of-day, and protocol fixed effects. Specifications with user fixed effects are estimated by computing the pre-post difference with user fixed effects in each year separately and then testing the null hypothesis that the difference-in-differences equals zero. The basic controls include trip duration, rain, temperature, and a weekend dummy. The full set adds distance to nearest hotspot and number of observers. Standard errors are robust to heteroskedasticity.

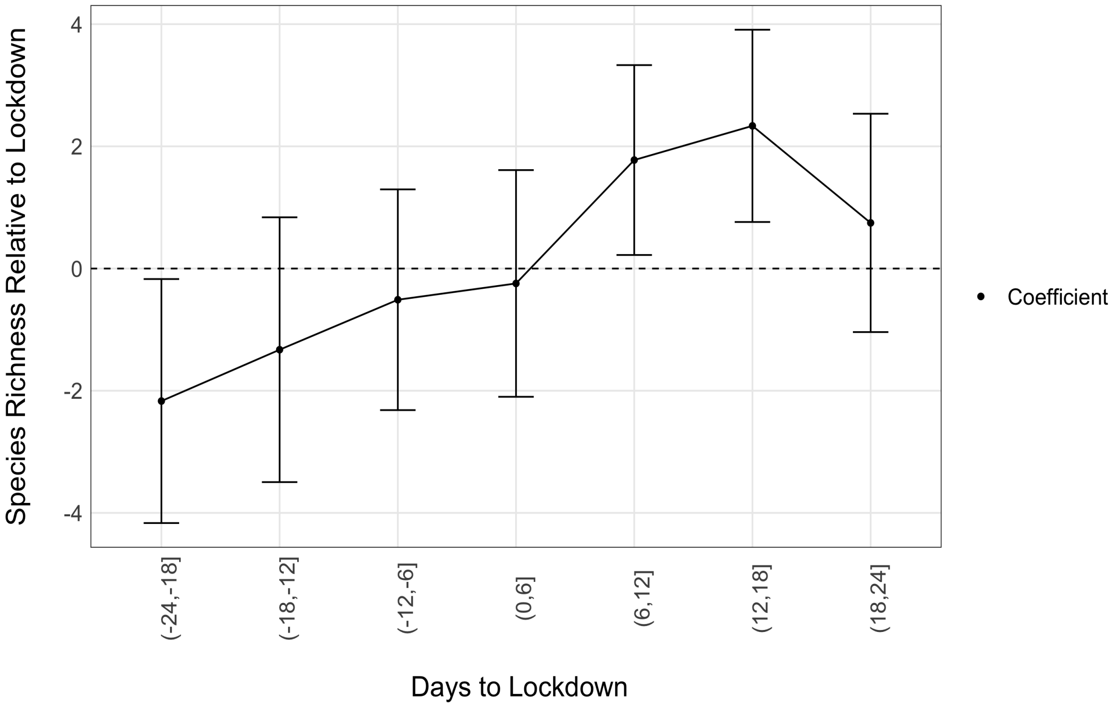

Figure S4: Dynamic Impact of Lockdown on Species Richness

Note: The x-axis denotes 6-day time bins. Negative values denote days before lockdown. The coefficient is the DD estimate for the impact of lockdown on species richness during a given time bin relative to that same bin in 2019. The period (-6,0] is omitted so all coefficients are relative to the week before (and including the day of) lockdown. Bars are 95% confidence intervals. The estimation sample is pre-processed for representativity, and include users meeting the 5-trip participation constraint. Observations are from the top 20 cities by population density. Data from March 22nd, 2020 (the Janta Curfew) are dropped. All regressions include district, hour-of-day, and protocol fixed effects, and control for trip duration, rain, temperature, and a weekend dummy. Standard errors are robust to heteroskedasticity.

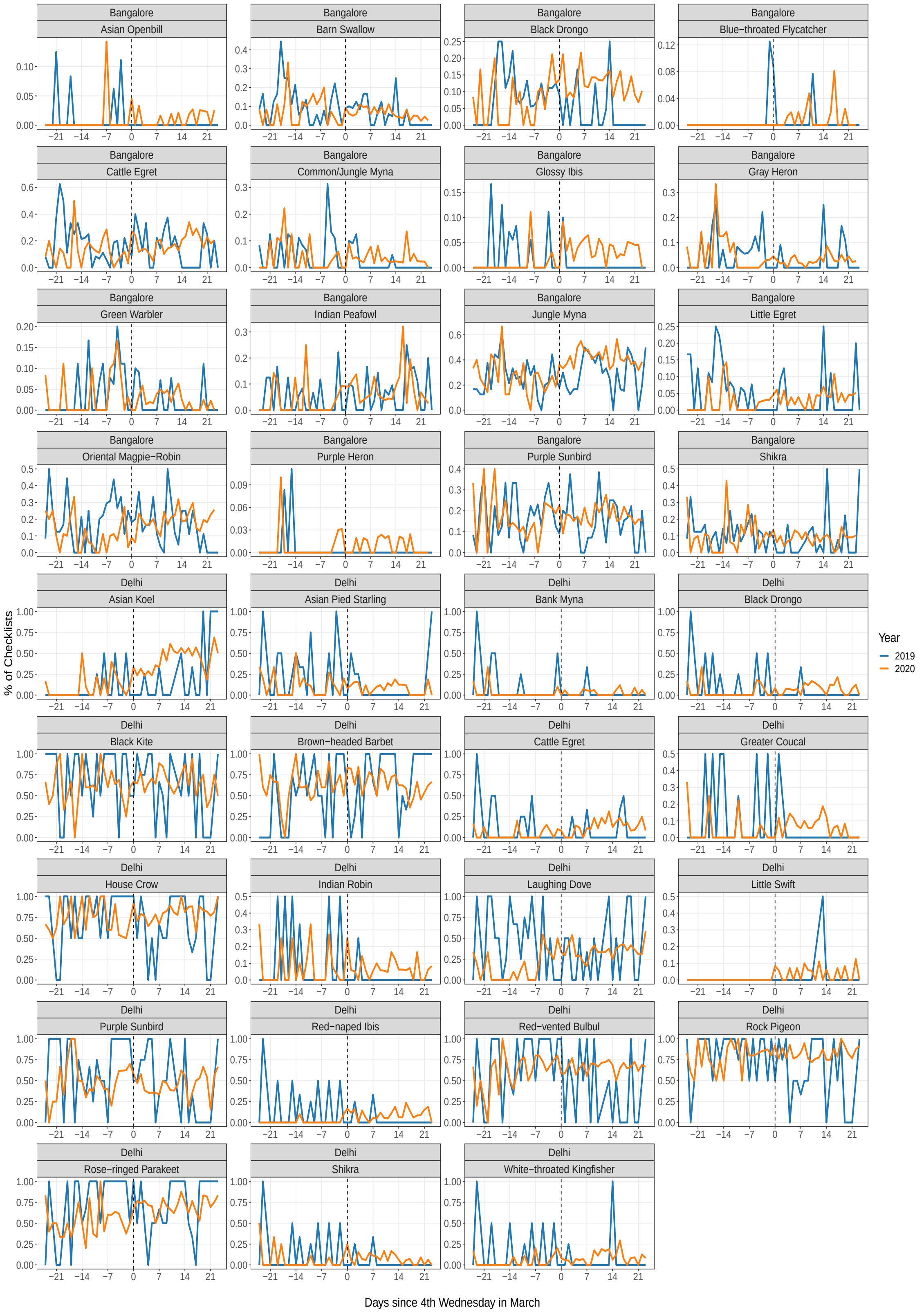

Figure S5: Daily Species Distributions (All Marginal Species)

Note: Species Distributions of marginal species in Delhi and Bangalore not shown in main text. Y-axis is the percentage of checklists reporting the species. X-axis is the number of days relative to the 4th Wednesday of March, which corresponds to the lockdown in 2020. The sample of checklists are stationary trips from the DD regression sample (2-trip participation constraint)

#### S2.2 Tables

Table S1: Breakup of Trip Protocols in Full Sample

| Protocol | Num. Trips | Pct. |
|---|---|---|
| Stationary | 48974 | 48.51 |
| Traveling | 44875 | 44.45 |
| Incidental | 6001 | 5.94 |
| Historical | 806 | 0.8 |
| Random | 161 | 0.16 |
| Area | 87 | 0.09 |
| Nocturnal Flight Call Count | 30 | 0.03 |
| Banding | 23 | 0.02 |
| eBird Pelagic Protocol | 4 | 0 |
| International Shorebird Survey (ISS) | 1 | 0 |

Note: The incidental protocol describes trips where birdwatching is not the primary purose. It could involve observing a species while cooking, or while driving. The historical protocol is for users who enter available data into eBird from their personal diaries, possibly going back decades. For the random protocol, a user travels 3 to 5 miles in a random direction from the last birding location. The area protocol is used when thoroughly searching a given area, and is often used by biologists when monitoring a specified site. The Nocturnal Flight Call is a community initiative to document nocturnally migrating birds. Banding involves using a net to capture, record, and release species. The eBird Pelagic Protocol is a specialized protocol for pelagic birding by boat. The ISS an international initiative to crowd-source information on wetland and shoreside birds.

Table S2: eBird Districts in the Top 20 by Population Density

| District | State |
|---|---|
| Bangalore | Karnataka |
| Central Delhi | Delhi |
| Chennai | Tamil Nadu |
| East Delhi | Delhi |
| Haora | West Bengal |
| Hyderabad | Telangana |
| Kolkata | West Bengal |
| Mumbai | Maharashtra |
| Mumbai Suburban | Maharashtra |
| New Delhi | Delhi |
| North 24 Parganas | West Bengal |
| North Delhi | Delhi |
| North East Delhi | Delhi |
| North West Delhi | Delhi |
| Rangareddy | Telangana |
| South 24 Parganas | West Bengal |
| South Delhi | Delhi |
| South West Delhi | Delhi |
| West Delhi | Delhi |

Note: The list describes districts in eBird that are a subset of the top 20 by population density. The full set is obtained by sorting a list of all 640 districts of India by population density and selecting the top 20. Among these top 20, eBird users recorded data in the 19 listed here.

Table S3: Birdwatching Activity Before and After Lockdown (2020)

| | Before | After |
|---|---|---|
| **Panel A: All Cities** | | |
| Num. Trips | 8561 | 14858 |
| Num. Cities | 269 | 167 |
| Num. Users | 542 | 542 |
| Num. Species | 1010 | 778 |
| **Panel B: Top 20** | | |
| Num. Trips | 1376 | 2699 |
| Num. Cities | 19 | 17 |
| Num. Users | 103 | 103 |
| Num. Species | 387 | 284 |

Note: Statistics are for 2020 only. "Before" corresponds to March 1st to March 24th 2020. "After" corresponds to March 25th to April 17th, 2020. Data include consistent users who recorded at least 2 trips before and after lockdown. Panel A include data from users across the country, and Panel B include users in districts/cities among the top 20 by population density. Statistics are aggregates over all users; for example, number of species is the total number of unique species collectively recording by all users.

Table S4: Trip Characteristics Before and After Lockdown (2020)

| | Before | After |
|---|---|---|
| **Panel A: All Cities** | | |
| Mean Trip Length | 37.72 | 25.91 |
| Mean Species Richness | 17.09 | 13.36 |
| Median Species Richness | 13 | 12 |
| Mean Dist. to Hotspot (km) | 2.39 | 1.9 |
| Mean Rainfall (mm) | 0.64 | 0.87 |
| Mean Temperature (Celsius) | 24.77 | 27.65 |
| **Panel B: Top 20** | | |
| Mean Trip Length | 36.85 | 25.71 |
| Mean Species Richness | 15.59 | 12.98 |
| Median Species Richness | 11 | 12 |
| Mean Dist. to Hotspot (km) | 0.8 | 0.81 |
| Mean Rainfall (mm) | 0.03 | 0.22 |
| Mean Temperature (Celsius) | 25.21 | 27.73 |

Note: Statistics are for 2020 only. "Before" corresponds to March 1st to March 24th 2020. "After" corresponds to March 25th to April 17th, 2020. Data include consistent users who recorded at least 2 trips before and after lockdown. Panel A include data from across the country, and Panel B includes users in districts/cities among the top 20 by population density. Statistics are on a per-trip basis. Distance to hotspot is calculated as the distance in kilometers from the trip's GPS coordinate to the GPS coordinate of the nearest hotspot as listed in the eBird database.

Table S5: Rarity of Marginal Species: Bangalore

| District | Species | Pct. | IUCN | Rarity |
|---|---|---|---|---|
| Bangalore | Jungle Myna | 29 | LC | common |
| Bangalore | Oriental Magpie-Robin | 16.91 | LC | common |
| Bangalore | Cattle Egret | 16.73 | LC | common |
| Bangalore | Purple Sunbird | 15.8 | LC | common |
| Bangalore | Shikra | 8.55 | LC | common |
| Bangalore | Barn Swallow | 7.62 | LC | common |
| Bangalore | Black Drongo | 6.32 | LC | common |
| Bangalore | Indian Peafowl | 5.39 | LC | common |
| Bangalore | Little Egret | 4.28 | LC | common |
| Bangalore | Common/Jungle Myna | 3.9 | | common |
| Bangalore | Gray Heron | 3.72 | LC | common |
| Bangalore | Green Warbler | 3.53 | LC | common |
| Bangalore | Black-headed Ibis | 2.42 | NT | common |
| Bangalore | Indian Scops-Owl | 1.86 | LC | common |
| Bangalore | Glossy Ibis | 1.49 | LC | common |
| Bangalore | Black-crowned Night-Heron | 1.12 | LC | rare |
| Bangalore | Asian Openbill | 0.74 | LC | rare |
| Bangalore | Blue-throated Flycatcher | 0.56 | LC | rare |
| Bangalore | Purple Heron | 0.37 | LC | rare |
| Bangalore | Black-rumped Flameback | 0 | LC | rare |

Note: Column (2) lists 19 marginal species identified through the algorithm described in section *Results*. Column (3) is the percent of total checklists (n=538) submitted during the 2019 study period reporting each marginal species. Rarity is classified on a global and local level. Column (4) documents the IUCN red list category (LC = Least Concern; NT = Near-Threatened) for each species obtained through BirdLife International. In column (5), species in the bottom 25th percentile of reporting frequency (column (3)) are locally rare Gaston (1994).

Table S6: Rarity of Marginal Species: Delhi

| District | Species | Pct. | IUCN | Rarity |
|---|---|---|---|---|
| Delhi | Rock Pigeon | 74.71 | LC | common |
| Delhi | House Crow | 72.41 | LC | common |
| Delhi | Rose-ringed Parakeet | 72.41 | LC | common |
| Delhi | Black Kite | 67.82 | LC | common |
| Delhi | Red-vented Bulbul | 62.07 | LC | common |
| Delhi | Brown-headed Barbet | 60.92 | LC | common |
| Delhi | Purple Sunbird | 55.17 | LC | common |
| Delhi | Laughing Dove | 42.53 | LC | common |
| Delhi | Coppersmith Barbet | 24.14 | LC | common |
| Delhi | Asian Pied Starling | 20.69 | LC | common |
| Delhi | Asian Koel | 12.64 | LC | common |
| Delhi | Red-naped Ibis | 12.64 | LC | common |
| Delhi | Cattle Egret | 11.49 | LC | common |
| Delhi | Greater Coucal | 11.49 | LC | common |
| Delhi | Shikra | 11.49 | LC | common |
| Delhi | White-throated Kingfisher | 10.34 | LC | common |
| Delhi | Alexandrine Parakeet | 9.2 | NT | common |
| Delhi | Black Drongo | 9.2 | LC | common |
| Delhi | Indian Robin | 9.2 | LC | common |
| Delhi | Black-rumped Flameback | 8.05 | LC | rare |
| Delhi | Bank Myna | 6.9 | LC | rare |
| Delhi | Large-billed Crow | 5.75 | LC | rare |
| Delhi | Little Swift | 1.15 | LC | rare |

Note: Column (2) lists 23 marginal species identified through the algorithm described in section *Results*. Column (3) is the percent of total checklists (n=87) submitted during the 2019 study period reporting each marginal species. Rarity is classified on a global and local level. Column (4) documents the IUCN red list category (LC = Least Concern; NT = Near-Threatened) for each species obtained through BirdLife International. In column (5), species in the bottom 25th percentile of reporting frequency (column (3)) are locally rare Gaston (1994).
